# FastEBM: Fast, Scalable, and Uncertainty-Aware Event-Based Disease Progression Modeling

**DOI:** 10.64898/2026.07.18.739221

**Authors:** Shayan Javid, Talia M. Nir, Alyssa H. Zhu, Ravi R. Bhatt, Leon M. Aksman, Neda Jahanshad, Alzheimer’s Disease Neuroimaging Initiative

## Abstract

Event-based models (EBMs) are used to infer ordering of biomarker alteration patterns with respect to disease progression. However, EBM approaches rely on computationally expensive permutation-based inference, assumptions of feature independence, and likelihood optimization that can limit scalability and stability in high-dimensional settings. Here, we introduce Fast Event-Based Model (FastEBM), a scalable, uncertainty aware, Markov-chain-based framework that reformulates disease progression inference as a subject-ordering problem on a data-driven diffusion manifold. The progression uncertainty, used to derive positional variance diagrams, is quantified using first-passage-time variability derived directly from the inferred Markov process.

Using synthetic experiments varying feature dimensionality, cohort size, noise level, and feature-correlation structure, we compared FastEBM with established methods, including Gaussian mixture model EBM (GMM-EBM), kernel density estimation EBM (KDE-EBM), and discriminative EBM (DEBM). FastEBM achieved the best accuracy and runtime. In low-subject/high-dimensional stress tests, FastEBM retained event-order recovery. FastEBM remained robust in simulations containing correlated and redundant features after decorrelation and feature-group handling. We applied FastEBM to real-world data to characterize biomarker progression in Alzheimer’s disease. First, we evaluated a low-dimensional multi- modal dataset from The Alzheimer’s Disease Prediction Of Longitudinal Evolution (TAD- POLE) challenge. Second, to demonstrate high-dimensional disease progression mapping, we applied FastEBM to regional cortical tau-PET data from the Alzheimer’s Disease Neuroimaging Initiative (ADNI). In both cases, FastEBM recovered progression patterns broadly consistent with the literature, also revealing lateralized progression trends.

These results show that diffusion-based Markov geometry provides a scalable and robust alternative to conventional event-based modeling. FastEBM is available at: https://github.com/sjusc07/FastEBM.

## 1 Introduction

Biological changes associated with neurodegenerative disorders, such as Alzheimer’s disease, often progress over decades before the emergence of clinical symptoms. This progression can be reflected in gradual changes across multiple biomarkers, such as neuroimaging measures, blood- and cerebrospinal fluid (CSF)-based biomarkers, and cognitive scores. Characterizing the temporal ordering of these biomarker changes is critical for understanding mechanisms of disease progression, enabling early interventions, stratifying individuals by disease stage, and informing clinical trial design (Young et al., 2024). However, reconstructing long-term disease trajectories remains challenging due to the limited availability of densely sampled longitudinal data and the difficulty of identifying individuals prior to symptom onset (Donohue et al., 2014; Hampel et al., 2018; Oxtoby & Alexander, 2017). As a result, many studies rely on cross-sectional or sparsely-sampled longitudinal datasets, necessitating computational approaches that can infer temporal structure from incomplete observations.

Data-driven disease progression models have emerged as a powerful framework for characterizing disease evolution directly from patient data, providing an alternative to traditional stage-based disease classifications (Ganjgahi et al., 2025). Among these methods, event-based models (EBMs) (Fonteijn et al., 2012) map disease progression as a sequence of discrete biomarker abnormality events without requiring *a priori* clinical staging. In this framework, each biomarker transitions from a normal state to an abnormal state, allowing estimation of the most likely ordering of these biomarker events across a population from cross-sectional data. Classical EBM approaches successfully infer this ordering by using probabilistic models of biomarker distributions together with likelihood-based optimization or sampling over permutations of possible event sequences. While these foundational methods have advanced the study of neurodegenerative diseases, scaling them introduces a trade-off between computational tractability, model complexity, and robustness. First, while searching across event permutations ensures a thorough exploration of the event space, the combinatorial approach leads to substantial computational cost as the number of features increases. Second, while assuming feature independence and using simplified distributional forms (Oxtoby, 2023), keeps the models mathematically practical and widely applicable, such frameworks are often constrained when applied to neuroimaging data where features vary widely in their underlying parametric shapes (Firth et al., 2020) and have intrinsic correlations that violate standard independence assumptions (Fonteijn et al., 2012; Venkatraghavan et al., 2019). Third, although likelihood-based and sampling-based procedures have improved the characterization of disease progression, their inferred orderings can remain sensitive to noise in the underlying feature measurements, potentially reducing sequence stability and reproducibility (Venkatraghavan et al., 2019).

In this work, we introduce Fast Event-Based Model (FastEBM). FastEBM addresses these limitations by reformulating disease progression modeling as a problem of subject ordering on a data-driven manifold (Coifman & Lafon, 2006). Rather than estimating a global permutation of biomarker events directly, FastEBM constructs a similarity graph across subjects and models disease progression as a diffusion process on this graph. A Markov chain defined on the graph captures the intrinsic geometry of the data and the corresponding fundamental matrix (Kemeny & Snell, 1969), provides a global representation of subject relationships designed to improve robustness to noise. Subjects are then ordered along a continuous progression axis relative to a control-derived reference, enabling reconstruction of biomarker trajectories and inference of event orderings via change point detection (Truong et al., 2020).

FastEBM addresses several limitations of existing event-based models. First, it does not require all predefined diagnostic or clinical stage labels, such as cognitively normal (CN), mild cognitive impairment (MCI), and Alzheimer’s disease (AD), prior to model fitting. Instead it defines biomarker abnormality through normalization relative to a control population, allowing progression to be inferred directly from the data. Although not a requirement, when such labels are available, they can be used to help define the disease trajectory. Second, FastEBM handles correlated biomarkers by combining subject-level modeling with an explicit preprocessing step that groups correlated features using hierarchical clustering and reduces redundancy using principal component analysis. This ensures that relationships between subjects reflect underlying biological variation rather than redundant feature contributions. Third, we implement an uncertainty-aware extension of FastEBM that characterizes uncertainty in the inferred progression ordering using first-passage-time variability derived from the same Markov process used to define disease progression. Unlike conventional uncertainty estimation approaches based on bootstrap resampling or permutation sampling, this framework derives uncertainty estimates directly from the intrinsic stochastic geometry of the inferred progression manifold. Finally, by avoiding combinatorial optimization over event permutations, FastEBM achieves substantial improvements in computational efficiency and scalability, making it suitable for high-dimensional datasets while preserving interpretability in the original biomarker space.

Here, we first evaluate a deterministic and an uncertainty-aware FastEBM using a comprehensive simulation framework designed to assess accuracy, computational efficiency, robustness to noise, and sensitivity to correlated features. Comparisons are performed against established EBM variants, including Gaussian Mixture Model (GMM)-EBM (Young et al., 2014), Kernel Density Estimation (KDE)-EBM (Firth et al., 2020), and discriminative EBM (DEBM) (Venkatraghavan et al., 2019). Next, we apply the framework to data from The Alzheimer’s Disease Prediction Of Longitudinal Evolution (TADPOLE) challenge (Marinescu et al., 2019) previously used to test other EBM models. Finally, we highlight FastEBM’s high-dimensional capabilities by applying it to localized regional tau-PET data from the Alzheimer’s Disease Neuroimaging Initiative (ADNI) to assess the biological plausibility of the inferred progression trajectories, and to map the estimated progression of tau-PET aggregation across more fine-grained cortical gyri rather than larger lobular regions. Throughout the remainder of the paper, unless otherwise specified, the uncertainty-aware version of FastEBM is referred to simply as FastEBM, while the version without uncertainty quantification is referred to as Deterministic FastEBM. The overall pipeline and experiments are freely available as an open source Python package (FastEBM; https://github.com/sjusc07/FastEBM).

## 2 Methods

### 2.1 FastEBM Modeling

FastEBM represents disease progression as a stochastic diffusion process over a subject similarity graph and infers feature event ordering from the resulting Markov chain progression geometry, as seen in **Figure** 1. The following subsections describe each stage of the modeling pipeline, including data representation, subject ordering, event-order inference, and uncertainty quantification.

**Figure 1:**
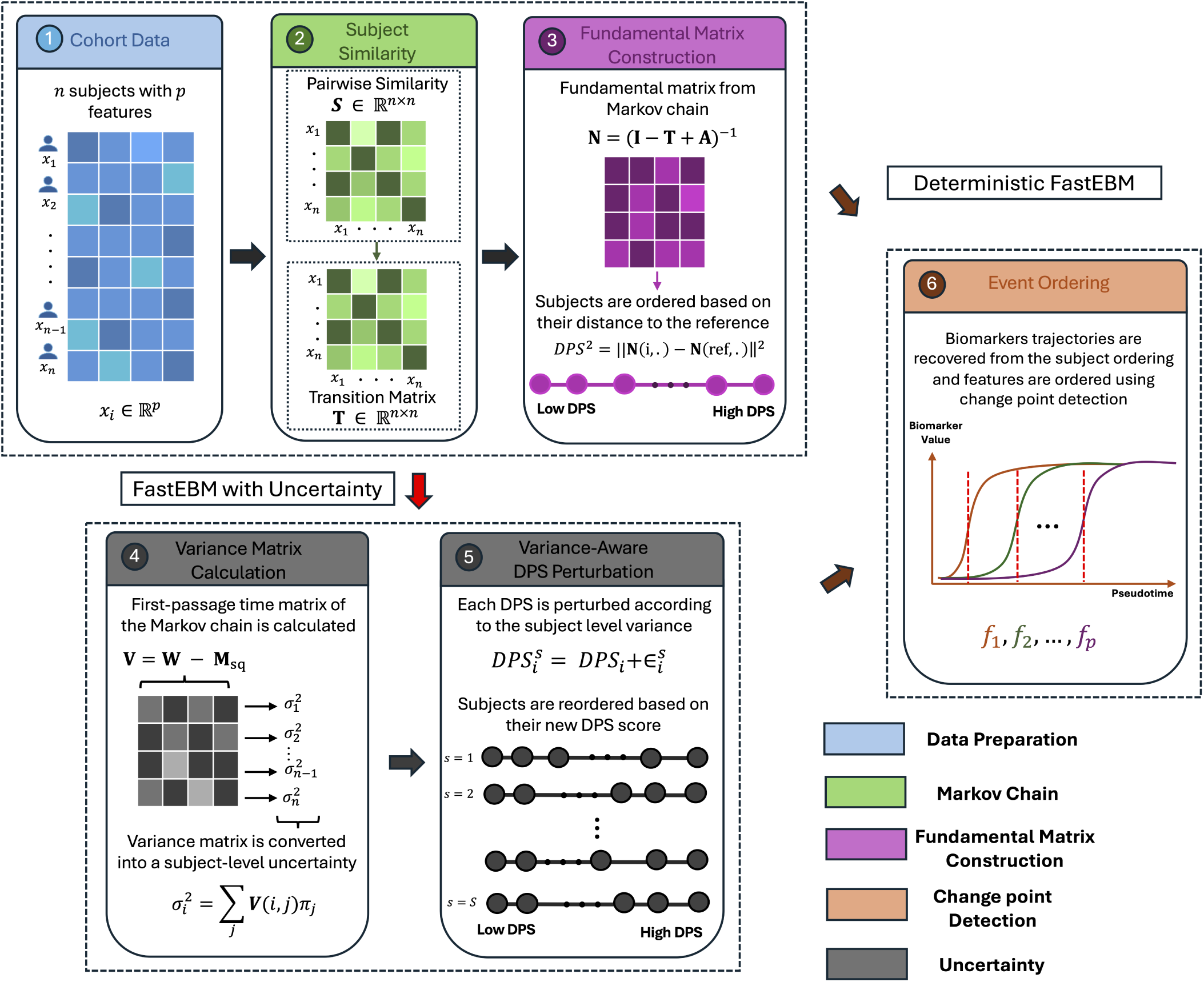
Overview of FastEBM with intrinsic uncertainty quantification. 1) Data are z-scored relative to a reference group. 2) Similarities between subjects are calculated using a locally-weighted Gaussian kernel. A Markov chain is constructed on the similarity-based transition matrix. 3) The fundamental matrix of the Markov chain is calculated and used to order subjects based on their distance to the reference template. 4) To quantify uncertainty in FastEBM, the first-passage variance matrix of the Markov chain is calculated and is converted to subject-level uncertainty. 5) Each disease progression score (DPS) is perturbed for a total of *S* iterations. 6) Feature trajectories are recovered from ordered subjects. Features are then ordered using a change point detection ordering. Deterministic FastEBM does change point detection directly after fundamental matrix calculation.

#### 2.1.1 Data Representation

Let Ω = {*x*_1_, · · · *, x_n_*} denote a set of *n* subjects, each represented by a *p*-dimensional feature vector (*x_i_* ∈ R*^p^*). Partially missing data are handled using a weighted Euclidean distance formulation (see *Subject Similarity Graph Construction* section). All features are then z-scored relative to a user-defined control group

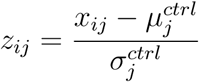

Where 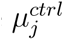and 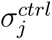 are the mean and standard deviation of feature *j* in the control group. This transformation aligns all features onto a common scale, encodes deviation from normative control (healthy) values, and allows disease progression to be inferred without explicit stage labels.

#### 2.1.2 Subject Similarity Graph Construction

FastEBM starts by constructing a weighted similarity matrix, **S** ∈ R*^n×n^* using a locally adaptive Gaussian kernel (Zelnik-Manor & Perona, 2004):

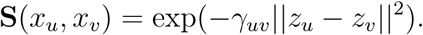

Where ||*z_u_* − *z_v_*|| is the Euclidean distance in the normalized space. The kernel bandwidth, *γ_uv_* is determined using local neighborhood scaling defined as:

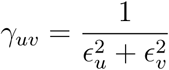

where *ɛ_u_* denotes subject *x_u_*’s distance to its *k*-th nearest neighbor. In the presence of partially missing values, we used a weighted Euclidean distance (Dixon, 1979) defined as

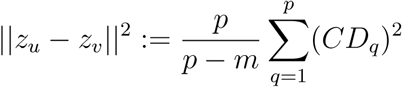

where *m* is the maximum number of missing values in either vector. *CD_q_* for the *q*-th feature will be zero if either subject is missing that feature, otherwise it is defined as the difference between the feature values (*z_uq_* −*z_vq_*). To correct for sampling density bias, we followed Coifman and Lafon, 2006 and applied kernel normalization

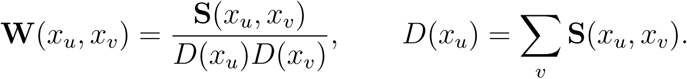

Finally, using row normalization (Chung, 1997) we can define a right-stochastic transition matrix **T**(*x_u_, x_v_*) as

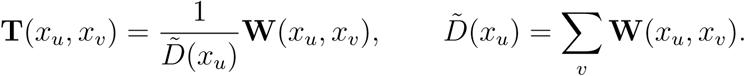

This defines a Markov chain over the subjects, where **T**(*x_u_, x_v_*) represents the probability of transitioning between subjects in the latent disease space. Because the graph is fully connected with nonzero transition probabilities, the chain is ergodic and has a unique stationary distribution (Kemeny & Snell, 1969).

#### 2.1.3 Fundamental Matrix Calculation

Because **T** is row-stochastic, it has a stationary distribution *π*, defined by

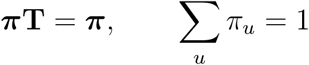

where *π_u_* denotes the stationary probability of subject *u*. We used this stationary distribution to form the rank-one stationary matrix

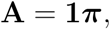

where **1** is a column vector of ones. Thus, each row of **A** is equal to the stationary distribution. For an ergodic finite Markov chain, **A** is also the limiting transition matrix, **A** = lim*_m→∞_* **T***^m^*, and represents the equilibrium component of the chain (Kemeny & Snell, 1969). To look at the global relationships between subjects, beyond nearest neighbors, we computed the fundamental matrix of the Markov chain (Kemeny & Snell, 1969):

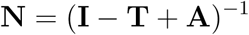

The fundamental matrix encodes the expected number of visits to each subject under the diffusion process. This provides a measure of reachability and connectivity within the graph. Subjects that lie along similar regions of disease trajectory will have higher mutual visitation frequencies, while subjects that are distant or belong to different regions of the manifold will have weaker connections. Each subject *x_i_* is represented by the corresponding row **N**(*i, .*), yielding a diffusion based representation that reflects its position in the global structure of the dataset.

#### 2.1.4 Disease Progression Axis

Following the diffusion-based embedding framework of Haghverdi and colleagues, (Haghverdi et al., 2016) we used the fundamental matrix as a new feature representation of the data and defined a disease progression axis by ordering the subjects according to their distance from the reference template in the fundamental matrix space:

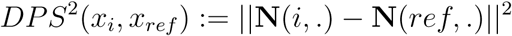

Here, the reference template, *x_ref_*, is defined as the median of the control group in the diffusion space. This provides an anchor corresponding to the normative feature profile. Subjects are then ranked according to their disease progression score (DPS) from this reference, producing a one-dimensional ordering that reflects increasing deviation from control states. This ordering is interpreted as the progression of disease stages.

#### 2.1.5 Feature Trajectory Reconstruction and Event Ordering

Feature trajectories are reconstructed by mapping each feature onto the inferred subject ordering. For each feature, values are sorted according to the subject rank. This approach, in theory, does not impose parametric assumptions or monotonicity constraints. Event ordering was estimated by identifying, for each feature, the position along the inferred disease-progression and selecting the position that maximizes the difference in mean feature value between subjects before and after the split. After subjects were ordered by their DPS, each feature was represented as a one-dimensional sequence

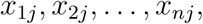

where *x_ij_*is the value of biomarker *j* for the *i*-th subject in DPS order.

For each feature, a single change point *τ_j_* was estimated by minimizing the within-segment sum of squared errors under a two-segment piecewise-constant model:

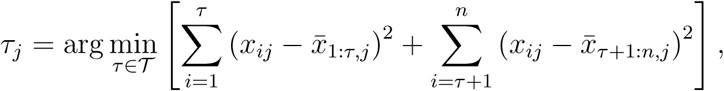

where *x̄*_1:*τ,j*_ and *x̄_τ_*_+1:*n,j*_ are the mean feature values before and after the candidate change point.

Candidate change points were constrained to satisfy a minimum segment-size requirement so that both pre-change and post-change segments contained at least two observations to prevent extreme single points from affecting the change point location. The estimated event position for feature *j* is the disease-progression position corresponding to its minimizing change point *τ_j_*. Features are ordered by these estimated change point positions.

#### 2.1.6 Uncertainty Quantification

We developed an intrinsic Markovian uncertainty quantification procedure for FastEBM that propagates uncertainty from the geometry of the subject-level Markov chain to feature event ordering. Our method does not rely on subject resampling or bootstrap aggregation. Instead, it uses first-passage-time variability of the Markov chain to quantify how uncertain each subject’s disease progression score is with respect to the inferred disease-progression manifold. To quantify uncertainty in the DPS, we used the Kemeny-Snell first-passage-time variance matrix from the FastEBM Markov chain described above. For each subject, this variance describes how variable the number of Markov-chain steps is from that subject to other subject states. This is motivated by the interpretation of disease progression as movement over the subject manifold. If a subject has highly variable first-passage times to other states, then its location along the inferred progression axis is less stable. Conversely, if the chain reaches other states from a subject in a more predictable manner, that subject’s location is more stable. The mean first-passage time (Kemeny & Snell, 1969) in matrix form is calculated as

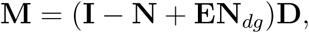

where **E** is a matrix of ones,

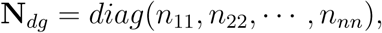

and

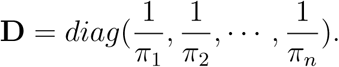

Then, the second moment matrix of first-passage is calculated using the Kemeny and Snell, 1969 expression

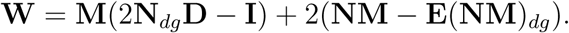

The first-passage variance matrix is then

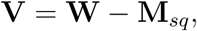

where

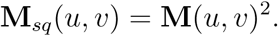

Thus, **V**(*u, v*) is the variance of the first-passage time from subject *u* to subject *v*. Where **V**(*u, v*) denotes the variance of the estimate of the number of steps required to reach state *v* starting from state *u*. To quantify the localized Markovian uncertainty for each subject, similar to Le et al., 2026, we compute a volatility metric but at the subject-level. The first-passage variance matrix is converted into a subject-level uncertainty score by averaging each subject’s first-passage variance over destination states weighted by the stationary distribution:

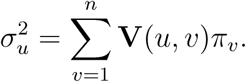

This aggregation gives more weight to states that are more representative of the long-run Markov chain and less weight to rarely visited states. The resulting *σ*^2^ is interpreted as the intrinsic Markovian uncertainty associated with subject *u*’s position on the disease-progression manifold.

Because *σ*^2^ is measured in Markov-chain time units rather than DPS units, we convert it into a relative uncertainty score by dividing each *σ*^2^ by the median of all the subject-level variances:

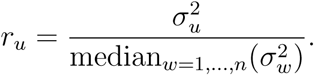

The quantity *r_u_* preserves relative variability across subjects while removing the arbitrary scale of the first-passage-time variance. To place the perturbation on the DPS scale without introducing a manually tuned global noise parameter, we estimate a data-derived local DPS resolution. Let

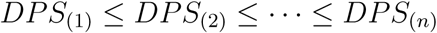

be the sorted baseline normalized DPS values. We define

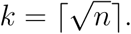

This choice gives a neighborhood size that increases with sample size while satisfying *k/n* → 0, a standard condition in nearest-neighbor nonparametric estimation (Stone, 1977). After sorting subjects by their baseline DPS values we computed the local-neighborhood DPS spacing as

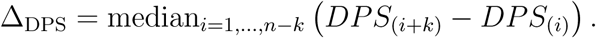

This quantity estimates the typical width of a local neighborhood along the fitted progression axis. The use of local neighbor distances as data-adaptive scale parameters follows the same principle as local scaling and adaptive bandwidth methods, where distances to nearby observations are used instead of a fixed global scale (Abramson, 1982; Zelnik-Manor & Perona, 2004). We used the median of the local spacings to obtain a robust scale estimate, reducing sensitivity to unusually large gaps or local irregularities in the DPS distribution (Huber, 2025).

The DPS noise variance for subject *u* is then

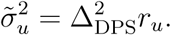

The squared scaling factor is required because variances transform quadratically under changes of scale. Thus, Δ_DPS_ determines the typical standard-deviation scale of the perturbation, while *r_u_* modulates this scale according to the subject-specific Markov first-passage variability. For each Monte Carlo replicate *s* = 1*, . . ., S*, we sample subject-level DPS noise

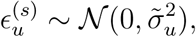

and perturb the baseline DPS:

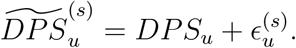

The perturbed DPS values are normalized to [0, 1], and subjects are reordered according 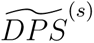. Feature event ordering is recomputed for each replicate using the same change point detection procedure as FastEBM. Across replicates, this produces a distribution of event positions and event ranks for each feature. An adaptive convergence criterion was used to determine the required number of perturbation samples. In this mode, the requested number of samples *S* was treated as an upper bound. After every batch of *s* samples, once at least *S_min_* samples had been generated, convergence was assessed. We looked at the convergence of mean feature event ranks. Let *r*^(^*^t^*^)^ denote the event rank of feature *k* in Monte Carlo sample *s*, using all samples generated up to checkpoint *t*. The mean event rank at checkpoint *t* was

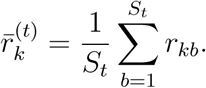

The maximum absolute change from the previous checkpoint was

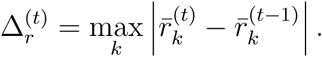

Sampling was stopped early when the selected convergence statistics remained below their specified tolerances for a required number of consecutive checkpoints:

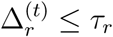

The pipeline for deterministic and uncertainty-aware FastEBM is shown in **Figure** 1.

#### 2.1.7 Correlated Feature Handling

To reduce the influence of highly correlated features on the inferred disease progression sequence, FastEBM includes an optional preprocessing step for detecting and merging correlated features. Correlated features were identified using pairwise feature correlations calculated using the feature matrix. A symmetric Pearson correlation matrix, **R**, was first estimated across features, and converted into a distance matrix **L**

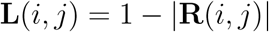

Hierarchical clustering with average linkage (Müllner, 2011) was then applied to **L**. Candidate feature clusters were then evaluated over a range of clustering thresholds. Singleton clusters were ignored. For each candidate cluster containing at least two features, the average within-cluster Pearson correlation was computed by first transforming each correlation value using Fisher’s *z* transformation (Corey et al., 1998)

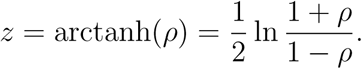

Then the average, 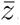, is calculated in *z*-space and transformed back to correlation scale using

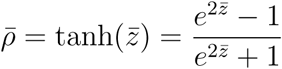

Clusters were retained only if their average correlation exceeded a user-specified threshold, set to 0.8 by default. After correlated clusters were identified, each cluster was merged into a single feature. For each correlated feature group, the original feature values were standardized, and principal component analysis was applied with one component. The first principal component was then used as a representative summary feature for that cluster. Features not assigned to any correlated cluster followed the regular FastEBM preprocessing pipeline. The resulting decorrelated dataset therefore contained all non-correlated original features with one principal-component-derived feature for each correlated cluster.

### 2.2 Experiments

We evaluated FastEBM in three experimental settings designed to assess: (i) comparative performance against existing event-based modeling methods, (ii) robustness to correlated and high-dimensional feature structure, and (iii) performance on real-world dataset. Synthetic experiments enabled direct comparison against known ground-truth event orderings, while real-world data experiments evaluated practical applicability towards mapping Alzheimer’s disease progression.

#### 2.2.1 Synthetic Data

We generated synthetic datasets to emulate disease progression using sigmoidal biomarker trajectories (Young et al., 2015). For each feature j, the trajectory was defined as:

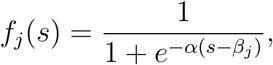

where *s* ∈ [0, 1] represents latent disease stage, *β_j_* defines the onset of abnormality, and *α* controls the slope of progression (set to 1). The parameters *β_j_* were uniformly spaced across features to define a known ground-truth ordering. Subjects were sampled uniformly along the disease axis, and feature values were perturbed by additive Gaussian noise:

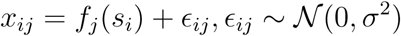

Control subjects were generated independently from N (0*, σ*^2^), resulting in datasets with equal numbers of control and diseased individuals. All experiments were conducted across the following set of configurations: we simulated datasets with different numbers of features, different numbers of subjects, and noise levels (**Table** 1). We varied one parameter at a time, resulting in 48 total synthetic datasets for the first two experiments. For experiment 1, we performed an additional stress-test and compared methods in a low-subject/high-dimensional setting, resulting in an additional 27 synthetic datasets (**Table** 1).

**Table 1:** Synthetic Data Experimental Configurations.

| Analysis | Number of Features | Sample Size | Noise | Configurations |
| --- | --- | --- | --- | --- |
| Main synthetic comparison | 10, 50, 100, 1000 | 500, 1000, 2000, 5000 | 0.2, 0.4, 0.8 | 48 |
| Low-subject/high-dimensional comparison | 100, 500, 1000 | 50, 100, 200 | 0.2, 0.4, 0.8 | 27 |

#### 2.2.2 Comparison of Methods

For our baseline comparisons of both simulated and real world data (Experiments 1 and 3), we evaluated FastEBM against three established event-based modeling approaches: 1) Gaussian mixture model (GMM) EBM (Young et al., 2014), which models biomarker distributions using Gaussian mixtures and infers event sequences using likelihood maximization; 2) kernel density estimation (KDE) EBM (Firth et al., 2020), which replaces Gaussian mixture modeling with KDE for greater flexibility; and 3) discriminative EBM (DEBM) (Venkatraghavan et al., 2019) which estimates subject-specific event orderings and aggregates them using a generalized Mallows model. For GMM-EBM and KDE-EBM we used the KDE-EBM package (https://github.com/ucl-pond/kde ebm) version 0.0.3. For DEBM we used the pyebm package (https://github.com/88vikram/pyebm) version 2.0.3.

#### 2.2.3 Evaluation Metrics

Kendall’s tau rank correlation coefficient (Kendall, 1938) was used to compare the inferred ordering to the ground truth ordering. Kendall’s tau measures the correspondence between two rankings. Tau values can range between -1 and 1 with values close to 1 indicating strong agreement and values close to -1 indicating strong disagreement. Runtime was measured for each method and for each dataset using Python’s time module. All experiments were run on an 2019 iMac desktop computer with a 3.9 GHz Intel Core i9 processor and 40GB of memory. For all the experiments we used a single core.

#### 2.2.4 Experiment 1: Performance Comparisons Across Methods

To evaluate the accuracy and computational efficiency of FastEBM relative to existing EBM approaches, we used the same dataset for all methods under each of the configurations listed in **Table** 1. For FastEBM, to maximize the sensitivity to recover fine structure of the manifold (van Dijk et al., 2018), we used one nearest neighbor. All features were z-scored relative to the control subjects before running FastEBM. In case of ties in the feature rankings, the features were randomly ordered. For the uncertainty parameter we used 1000 iterations of noise samples, the median-template ordering method, and one nearest neighbor. Convergence was checked every 25 samples after a minimum of 20 samples, with event-rank convergence tolerance set to 0.25. Features were ordered based on their median rank. Mean values were used to break ties in feature rankings. DEBM was run using pyebm.debm.fit, with no additional factors (Factors = []). Stage labels were mapped to CN for controls and AD for patients. For KDE-EBM and GMM-EBM, mixture models were fitted using fit_all_kde_models and fit_all_gmm_models, respectively, both with implement_fixed_controls = True. Event orderings were then estimated using the package’s Markov Chain Monte Carlo (MCMC) routine with plotting disabled and otherwise default parameters. The same synthetic data-generation procedure, event-order ground truth, model-fitting pipeline, and evaluation metrics used in the main benchmark were retained. Runtime was measured in seconds for each successful method/configuration run, and ordering accuracy was measured by Kendall’s tau between the inferred event order and the known ground-truth order. Failed runs occurred only for KDE-EBM and were caused by non-positive-definite covariance matrices during density estimation. These runs were retained in the completion-rate summary but were excluded from runtime and Kendall’s tau summaries because no valid ordering or runtime endpoint was produced. Also, the final high-dimensional configuration (1,000 features, 5,000 subjects, noise level 0.8) was omitted for DEBM, GMM-EBM, and KDE-EBM due to prohibitively long runtimes and because the scaling behavior of these methods had already been established in earlier configurations. FastEBM and Deterministic FastEBM were still evaluated on this setting.

#### 2.2.5 Experiment 2: FastEBM’s Robustness to Correlated Features

We evaluated FastEBM on synthetic data with correlated features. We used the same configurations to generate 48 datasets. For each configuration, we first generated exactly *f* features using the same synthetic data generation procedure described in section 2.2.1. For each dataset, 20% of the *f* base features were randomly selected as parent features for correlation augmentation. For each selected parent feature, 5 correlated copies were generated. Each copy was assigned its own correlation strength, sampled independently from a continuous range [0.2, 0.95]. Thus, correlation strength was not fixed at the dataset level, but was varied across copied features within the same dataset. Each parent feature was standardized (*parent_z_*), then for each sampled correlation strength for copy *k* (*ρ_k_*) we generated correlated copies using the Cholesky decomposition (Burgess, 2022) as

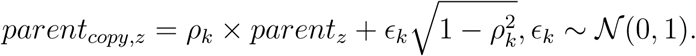

Each copy was then transformed back to the empirical mean and standard deviation of the parent feature. This construction yields an expected parent-copy correlation of approximately *ρ_k_*. Because correlated copies were intended to represent redundant measurements of the same latent event, no separate ground-truth disease-event order was defined among copies. We then used FastEBM’s correlated-feature workflow (described in section 2.1.7), to merge correlated/redundant features and obtain a decorrelated feature matrix to be used for FastEBM. Both the merged dataset and the original unmerged dataset were z-scored relative to the controls. FastEBM was then fitted on the merged, control-normalized dataset with 1 nearest neighbor. We used the same uncertainty parameters as experiment 1. The original unmerged data was then reordered according to the inferred subject pseudotime, and the final raw feature ordering was estimated based on the median of feature rankings with ties resolved using the mean ranking. This pipeline uses decorrelated data to derive subject ordering while preserving the original correlated features in the final feature ranking. To evaluate the performance of the FastEBM under these conditions, we computed the parent-level Kendall’s tau accuracy. The raw feature ordering was collapsed to a parent-level ordering by removing the correlated features from the final ordering. Then this ordering was compared with the true parent event ordering.

#### 2.2.6 Experiment 3: Method Comparison Using TADPOLE Data

As an assessment of biological validity in real-world data, we used the subset of the Alzheimer’s Disease Neuroimaging Initiative (ADNI) dataset used in The Alzheimer’s Disease Prediction Of Longitudinal Evolution (TADPOLE) challenge (Marinescu et al., 2019) to compare event ordering across EBM methods. We used subjects from ADNI 1, GO, and 2 available in the “D1 D2” dataset of the TADPOLE challenge (https://tadpole.grand-challenge.org). 15 subjects with missing intracranial volume (ICV) values were excluded. The final sample included 1722 participants: 519 cognitively normal (CN), 866 with mild cognitive impairment (MCI), and 337 with dementia (age range: 54.4-91.4; 44.8% female; **Table** 2). Following Venkatraghavan et al., 2019 and Du et al., 2023, we used 7 features: CSF amyloid-*β* 42 (ABETA42), total tau (TAU), and phosphorylated tau (PTAU); regional hippocampal and whole brain volumes; and Mini-Mental State Examination (MMSE) (Cockrell & Folstein, 2002), and the 13-item version of the Alzheimer’s Disease Assessment Scale Cognitive Subscale (ADAS13) (Mohs et al., 1997) cognitive assessments. We regressed out age and sex fitted on the CN group from cognitive scores, CSF measures, and brain volumes; ICV was also regressed out of brain volumes. Since each method has the capability of handling missing values for the features themselves, we did not remove any subjects with missing data in the 7 features. We used the GMM-EBM implementation in pyebm rather than the GMM implementation in KDE-EBM’s package for this experiment. The KDE-EBM implementation imputes missing feature values during probability-matrix construction. In the TADPOLE data, this step was numerically unstable for some features and prevented valid sequence inference. Thus, relative to experiment 1, the TADPOLE GMM-EBM results differs in software implementation and missing-data handling, while retaining the same modeling assumption of feature-wise Gaussian normal/abnormal distributions. For FastEBM all features were z-scored relative to the CN group. As before, for the uncertainty parameters we used 1000 iterations of noise samples, the same median-template ordering method, and one nearest neighbor. Sampling was done for 1000 iterations, convergence was checked every 25 samples after a minimum of 20 samples, with event-rank convergence tolerance set to 0.25. Features were ordered based on their median rank. MMSE, hippocampus, whole brain and ABETA42 decreased as stages progressed, thus, for EBM and KDE-EBM, we multiplied them by −1 to make these features increase over time. GMM and KDE methods were first fitted to the CN and dementia individuals, and the MCMC optimization step was done on the whole dataset (CN, MCI, and dementia).

**Table 2:** Demographic Data for TADPOLE.

|  | Total | CN | MCI | Dementia |
| --- | --- | --- | --- | --- |
| Participants, N | 1722 | 519 | 866 | 337 |
| Mean age (SD) | 73.8 (7.2) | 74.2 (5.8) | 73.0 (7.6) | 75.0 (7.8) |
| Female, N (%) | 772 (44.8%) | 267 (51.4%) | 354 (40.9%) | 151 (44.8%) |

#### 2.2.7 Experiment 4: Alzheimer’s Disease Cortical Tau PET Progression

To demonstrate the utility of FastEBM on higher dimensional data, we used FastEBM to evaluate regional cortical tau PET progression in Alzheimer’s Disease Neuroimaging Initiative (ADNI) participants. Flortaucipir (FTP) PET standardized uptake value ratio (SUVR) data for 64 bilateral Desikan-Killiany FreeSurfer cortical regions (Desikan et al., 2006) were downloaded from the ADNI database. The first available FTP PET scan of an individual with a diagnosis within six months of the scan was included in the analysis. We excluded scans that did not pass ADNI-provided quality control (QC). The final sample included 1,165 participants: 652 cognitively normal (CN), 389 with mild cognitive impairment (MCI), and 124 with dementia (age range 50.8-94.8; 54.5% female; **Table** 3). For each cortical SUVR, age- and sex-related effects were estimated using regression models fitted within the CN group and removed from all participants’ tau SUVR values. The resulting residuals were z-scored relative to the reference CN group before FastEBM fitting. FastEBM was then fitted to the 64 adjusted tau features using the median-template ordering method, one nearest neighbor, and an L2 change point model. Subject disease progression scores and the biomarker event ordering were estimated from the fitted model. Uncertainty in the inferred event ordering was quantified using FastEBM’s first-passage uncertainty procedure. The uncertainty analysis used the same parameter configurations as previous experiments.

**Table 3:** Demographic Data for ADNI Cortical Tau-PET.

|  | Total | CN | MCI | Dementia |
| --- | --- | --- | --- | --- |
| Participants, N | 1165 | 652 | 389 | 124 |
| Mean age (SD) | 72.6 (8.1) | 71.2 (7.8) | 73.7 (7.9) | 76.1 (8.8) |
| Female, N (%) | 634 (54.4%) | 412 (63.2%) | 173 (44.5%) | 49 (39.5%) |

## 3 Results

### 3.1 Performance Comparison Across Methods

Experiment 1 evaluated the runtime and ordering accuracy of five event-based modeling methods on synthetic datasets with varying numbers of features, cohort sizes, and noise levels. Ordering accuracy was quantified using Kendall’s tau against the known ground-truth event order, and computational cost was measured as wall-clock runtime in seconds. Because the low-subject/high-dimensional stress-test configurations were designed to probe a distinct failure mode, they were analyzed separately from the main benchmark. Across all configurations, KDE-EBM was the only method with failed runs due to the covariance matrix not being positive definite. These failures occurred in 36 configurations and were concentrated in higher-dimensional settings. Failures were most frequent at 1,000 features and at the lowest noise level. In the main benchmark, Deterministic FastEBM showed the strongest overall speed-accuracy profile. **Figure** 2 shows the Kendall’s tau accuracy of all the methods across the different configurations. Across all main-benchmark runs, Deterministic FastEBM achieved a median Kendall’s tau of 0.992 (IQR 0.934-1.000) with a median runtime of 1.28 seconds (s) (IQR 0.24-7.64 s). FastEBM achieved similarly high accuracy, with median Kendall’s tau 0.997 (IQR 0.946-1.000), but required longer runtime, with a median of 2.23 s (IQR 0.55-19.87 s). Thus, FastEBM preserved the event-ordering accuracy of Deterministic FastEBM while adding the expected computational cost of first-passage uncertainty estimation. The conventional EBM baselines were slower, less accurate, or less stable. DEBM achieved a median Kendall’s tau of 0.745 with a median runtime of 153.47 s. GMM-EBM achieved a median Kendall’s tau of 0.756 with a median runtime of 77.92 s, with a lower quartile accuracy of 0.127, indicating substantial degradation in a sizable fraction of the configurations. KDE-EBM performed the worst among the successful main-benchmark runs, with a median Kendall’s tau of -0.027 and a median runtime of 86.85 s; it also failed in 17 main-benchmark configurations. Restricting the main benchmark to the 30 configurations in which all five methods completed successfully gave the same qualitative result: Deterministic FastEBM achieved a median tau of 0.963 at 1.00 s, FastEBM achieved median tau 0.988 at 1.67 s, DEBM achieved a median tau of 0.613 at 70.28 s, GMM-EBM achieved a median tau of 0.710 at 33.48 s, and KDE-EBM achieved a median tau of -0.027 at 86.85 s. **Figure** 3 shows runtimes across different numbers of features and subjects. Noise level was the main factor influencing ordering accuracy. At a noise level of 0.2, Deterministic FastEBM and FastEBM were effectively perfect in the main benchmark; both resulted in a median Kendall’s tau of 1.000. At a noise level of 0.4, the median tau remained high for Deterministic FastEBM and FastEBM (0.992 and 0.997, respectively). At a noise level of 0.8, the accuracy decreased, but the FastEBM methods still outperformed the baselines: median tau was 0.903 for Deterministic FastEBM and 0.919 for FastEBM, compared with 0.333 for DEBM, 0.451 for GMM-EBM, and -0.252 for KDE-EBM. These results show that all methods were affected by severe noise, but FastEBM retained substantially more event-order information than the alternatives.

**Figure 2:**
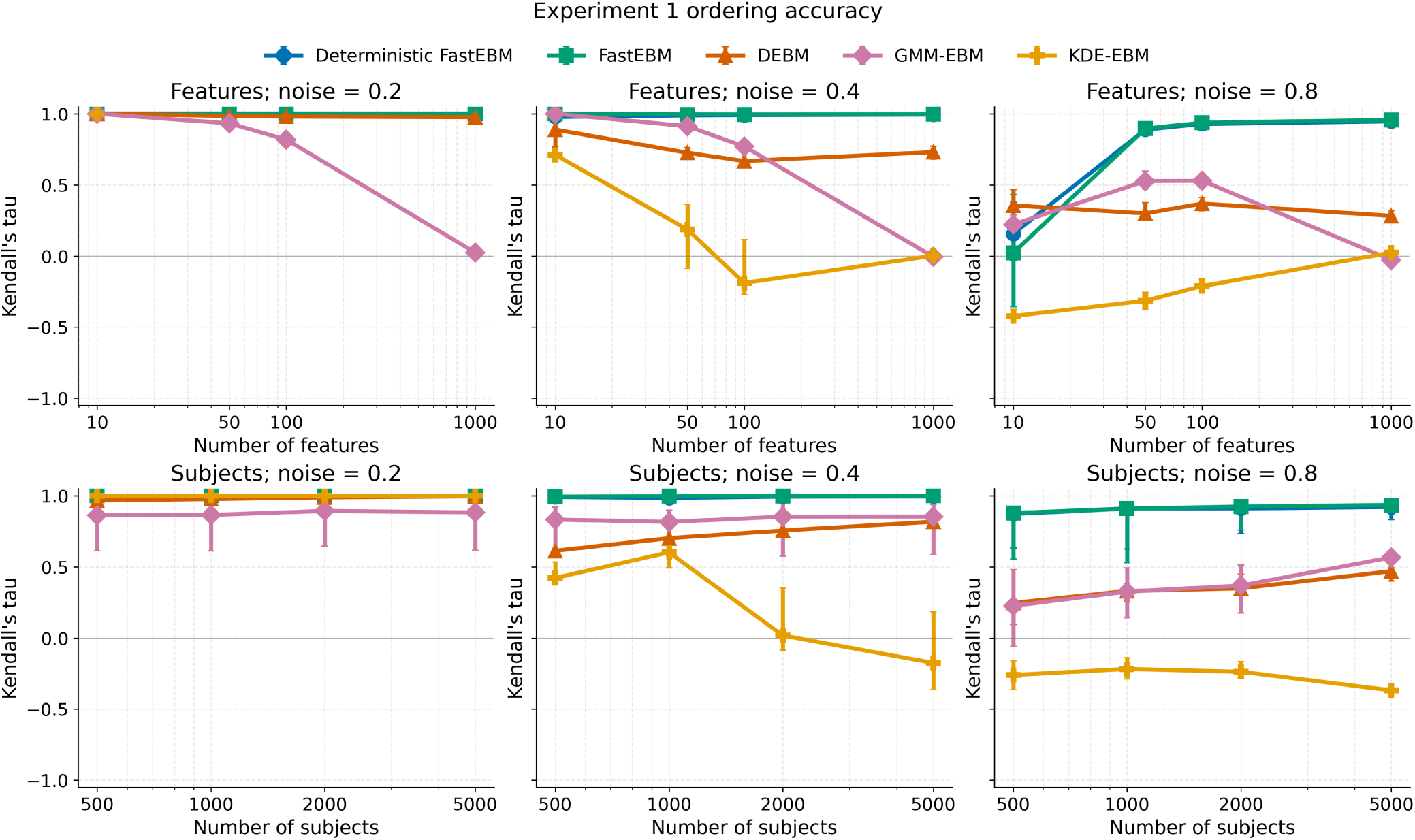
Kendall’s tau event-ordering accuracy in the main synthetic benchmark, excluding the high-dimensional low-sample-size configurations. Accuracy was computed by comparing the inferred event ordering with the true simulated ordering and is shown across feature counts, cohort sizes, and noise levels. FastEBM and Deterministic FastEBM achieved consistently high ordering accuracy, with the strongest degradation observed only at the highest noise level. DEBM and GMM-EBM were less stable across configurations, and KDE-EBM showed the weakest accuracy among successful runs. Points indicate medians across successful runs; error bars indicate the interquartile range.

**Figure 3:**
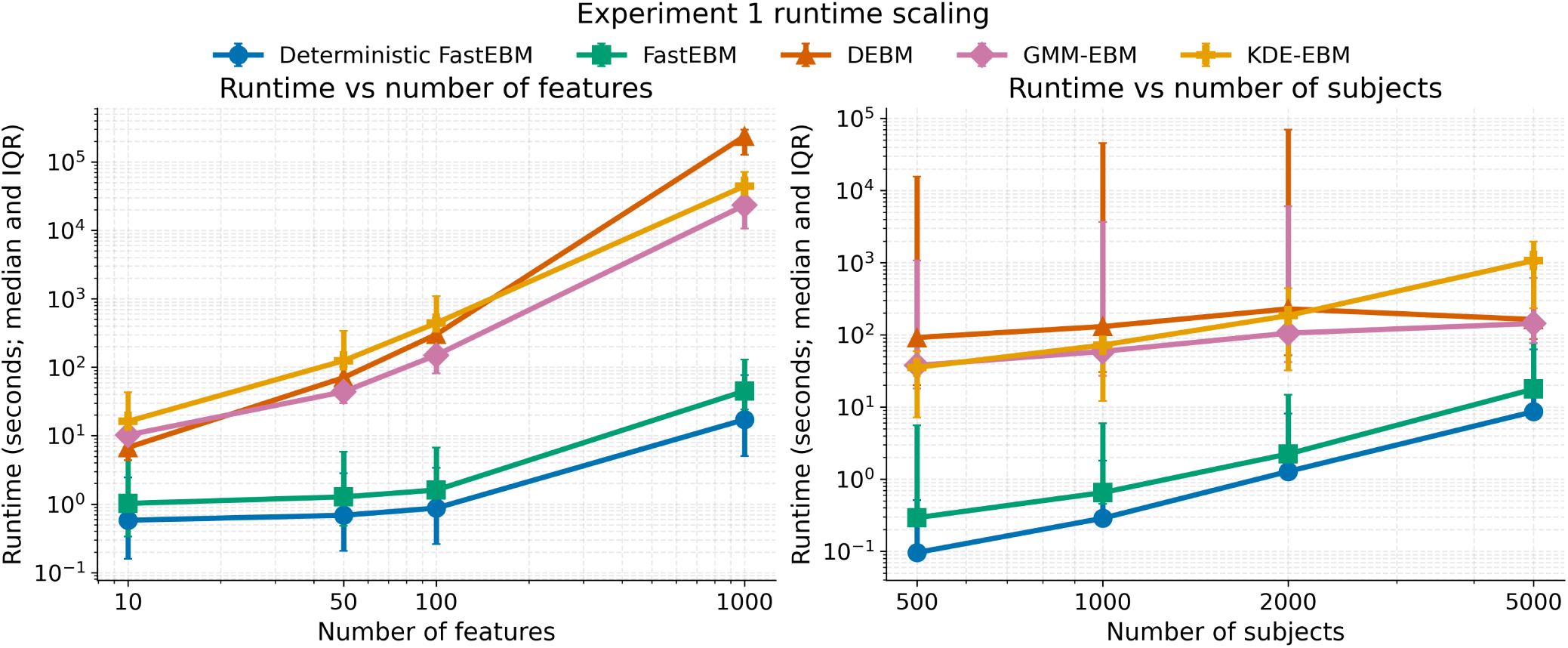
Runtime of Deterministic FastEBM, FastEBM, DEBM, GMM-EBM, and KDE-EBM across the main synthetic benchmark, excluding the high-dimensional low-sample-size configurations. Runtime is shown over total successful runs as a function of the number of biomarkers or cohort size. Deterministic FastEBM showed the lowest computational cost across the tested settings, while preserving scalability as feature count and sample size increased. FastEBM incurred additional runtime from the uncertainty estimations, and the conventional EBM baselines were substantially slower, particularly at larger feature counts. Points indicate medians across successful runs; error bars indicate the interquartile range.

To look at the scaling behavior, we highlight two representative experimental settings. At 1,000 total subjects, increasing the number of features from 10 to 1,000 increased Deterministic FastEBM’s median runtime from 0.19 s to 6.22 s while preserving high ordering accuracy, with median tau of 0.956 at 10 features and 0.994 at 1,000 features. In the same setting, DEBM runtime increased from 4.06 s to 235,951.53 s, GMM-EBM from 7.49 s to 22,995.34 s, KDE-EBM from 11.65 s to 11,644.34 s among successful runs, and FastEBM from 0.40 s to 38.91 s. At 100 features, increasing cohort size from 500 to 5,000 total subjects increased Deterministic FastEBM’s median runtime from 0.10 s to 9.32 s, while the median tau remained between 0.982 and 0.994. In contrast, the baseline methods remained slower and less accurate across this subject-scaling slice.

The low-subject/high-dimensional stress-test produced the same qualitative ranking but with more severe degradation of the baseline methods. In this subset, Deterministic FastEBM and FastEBM maintained high ordering accuracy, with median tau values of 0.965 and 0.962, respectively. Deterministic FastEBM remained extremely fast in this regime (median runtime 0.034 s), whereas FastEBM required 4.96 s. DEBM and GMM-EBM were substantially slower and less accurate, with median tau values of 0.445 and 0.011 and median runtimes of 1,925.48 s and 458.04 s, respectively. KDE-EBM completed only 8 of 27 low-subject/high-dimensional configurations and achieved median tau 0.059 among successful runs. **Table** 4 shows median Kendall’s tau and median runtimes across noise levels.

**Table 4:** Low-subject/High-dimensional Comparison Across Noise Levels.

| Noise Level | Method | Kendall's tau Median | Runtime Median (seconds) |
| --- | --- | --- | --- |
| 0.2 | DET-FastEBM<sup>a</sup> | 0.998 | 0.036 |
|  | FastEBM | 0.997 | 4.386 |
|  | GMM-EBM | 0.011 | 414.802 |
|  | KDE-EBM<sup>b</sup> | - | - |
|  | DEBM | 0.895 | 1022.935 |
| 0.4 | DET-FastEBM | 0.965 | 0.039 |
|  | FastEBM | 0.962 | 4.974 |
|  | GMM-EBM | 0.202 | 458.043 |
|  | KDE-EBM<sup>b</sup> | 0.484 | 49.331 |
|  | DEBM | 0.423 | 1053.824 |
| 0.8 | DET-FastEBM | 0.741 | 0.031 |
|  | FastEBM | 0.752 | 5.261 |
|  | GMM-EBM | -0.005 | 480.948 |
|  | KDE-EBM<sup>b</sup> | 0.031 | 49.047 |
|  | DEBM | 0.133 | 588.893 |

<sup>a</sup> Deterministic FastEBM.
<sup>b</sup> KDE-EBM failed for all runs with noise of 0.2, completed 3/9 runs for noise level 0.4, and 5/9 for 0.8.

Together, these results indicate that the low-subject/high-dimensional setting primarily exposes the instability and poor accuracy of the baseline approaches, while Deterministic FastEBM and FastEBM preserve useful event-order recovery at negligible computational cost. **Figure** 4 and **Figure** 5 show the runtime and accuracy across all methods for all the configurations.

**Figure 4:**
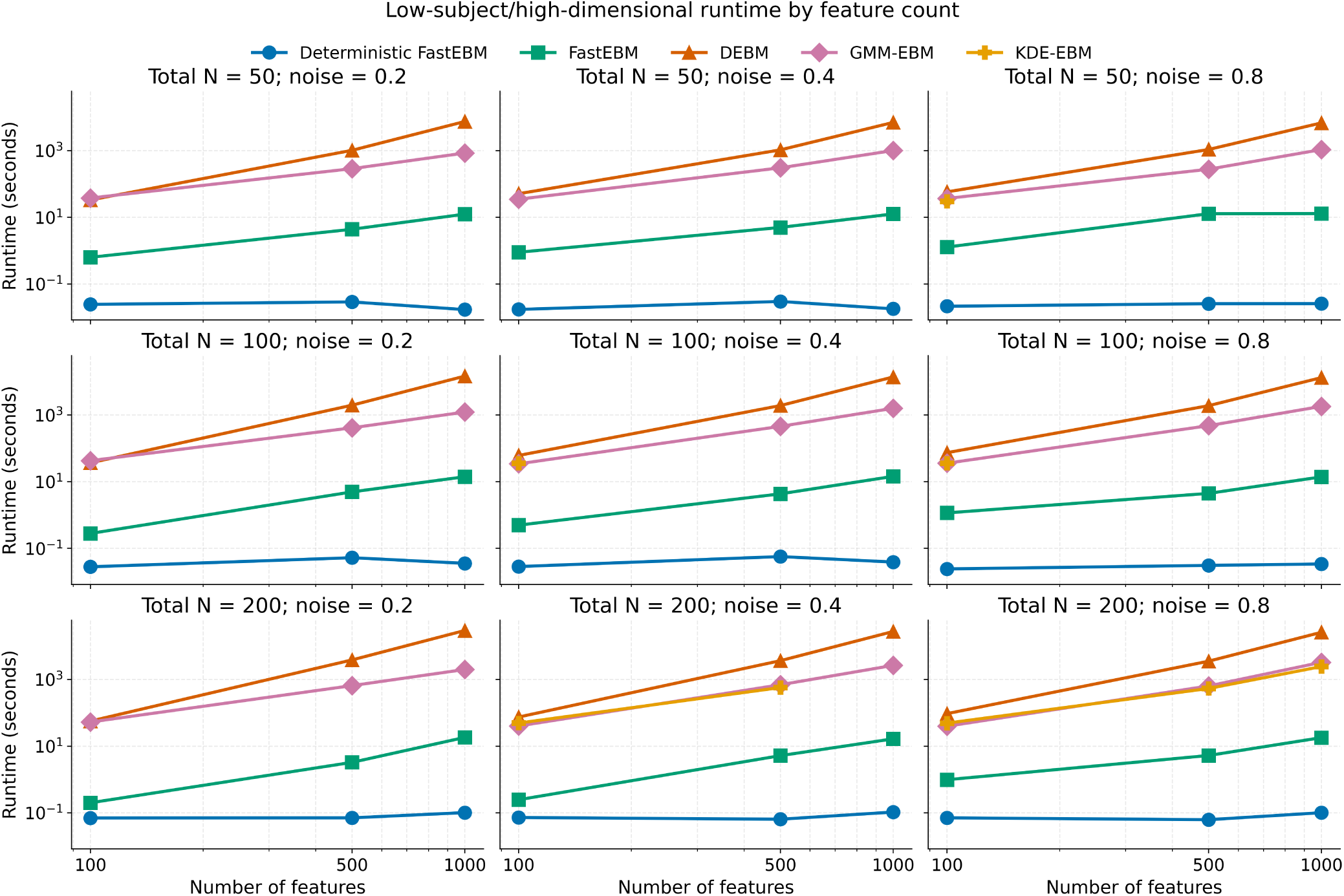
Runtime of Deterministic FastEBM, FastEBM, DEBM, GMM-EBM, and KDE-EBM in the high-dimensional low-sample-size synthetic benchmark, defined by 100-1,000 biomarkers, 25-100 subjects per group, and noise levels of 0.2, 0.4, and 0.8. Points and lines summarize successful runs across the high-dimensional low-sample-size configurations. FastEBM framework remained computationally lightweight across the stress-test settings, whereas DEBM and GMM-EBM required substantially longer runtimes. KDE-EBM completed only a subset of high-dimensional low-sample-size configurations.

**Figure 5:**
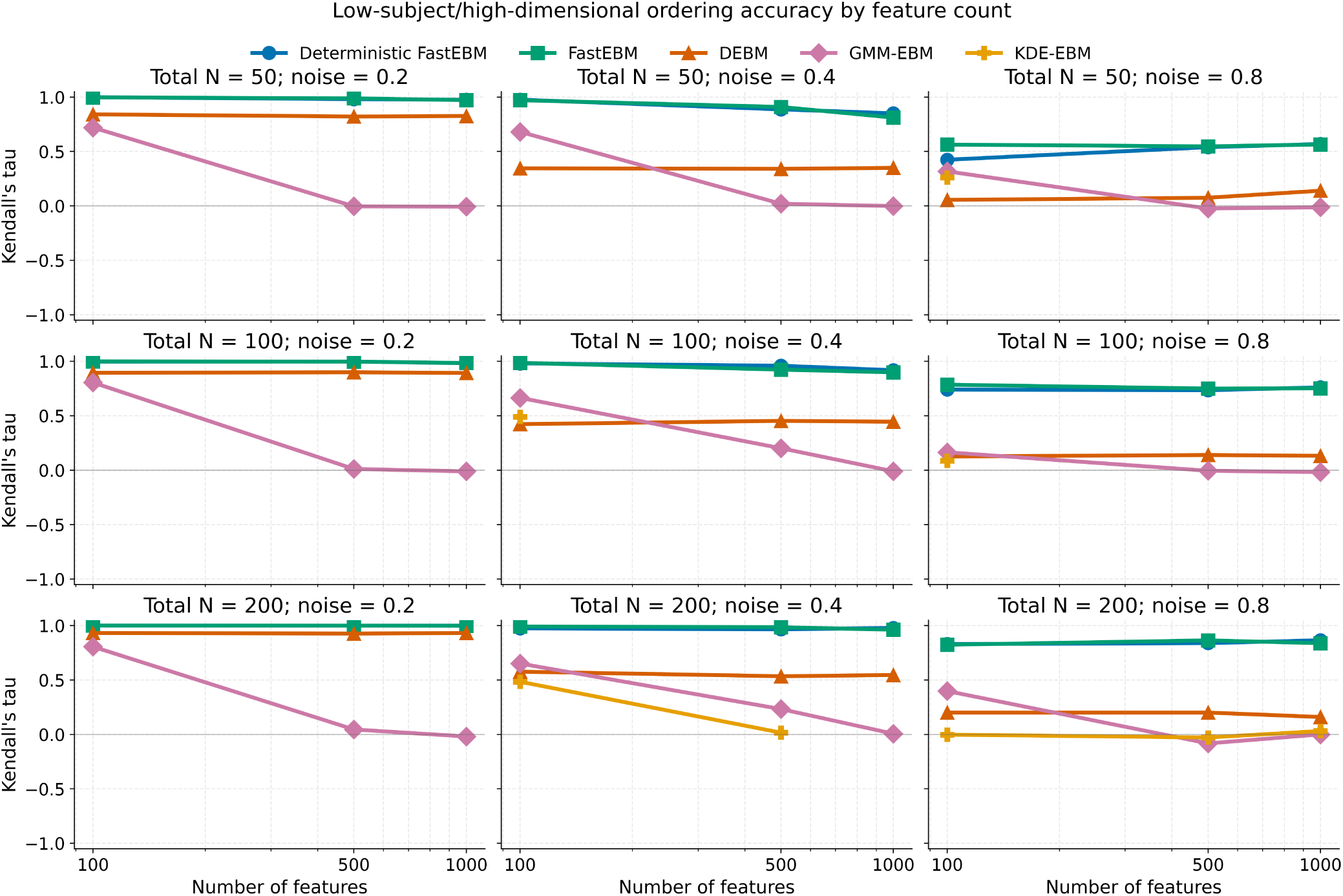
Kendall’s tau accuracy in the high-dimensional low-sample-size synthetic benchmark. Accuracy was computed by comparing the inferred event ordering with the true simulated ordering across high-dimensional low-sample-size configurations. Deterministic FastEBM and FastEBM retained high ordering accuracy despite the unfavorable ratio of features to subjects, while DEBM and GMM-EBM showed marked degradation. KDE-EBM had limited coverage because many high-dimensional low-sample-size configurations failed, and its successful runs showed substantially lower accuracy than the FastEBM- based methods.

### 3.2 Correlated Feature Robustness

When evaluating FastEBM in synthetic datasets containing correlated and redundant features, we generated datasets containing 10, 50, 100, or 1,000 base features, with 20% of base features selected as correlation parents and five correlated features generated per selected parent. This produced total feature counts of 20, 100, 200, and 2,000, respectively. FastEBM recovered the parent-level event ordering with high accuracy despite the presence of redundant correlated features. The mean parent-level Kendall’s tau was 0.858 and the median was 0.957 (IQR 0.864-0.987).

Noise was the dominant determinant of ordering accuracy. At noise level 0.2, parent-level ordering was near perfect, with a median Kendall’s tau of 0.994 (IQR 0.985-0.998). At noise level 0.4, performance remained high, with median tau equal to 0.959 (IQR 0.925-0.978). At noise level 0.8, accuracy decreased to a median tau of 0.788 (IQR 0.569-0.858), showing that severe measurement noise remained the primary failure mode even after decorrelating redundant features (**Figure** 6). The degradation at high noise was most noticeable in the smallest feature setting: with 10 base features, the median tau at noise 0.8 was 0.133, whereas the corresponding medians for 50, 100, and 1,000 base features were 0.776, 0.857, and 0.871, respectively.

**Figure 6:**
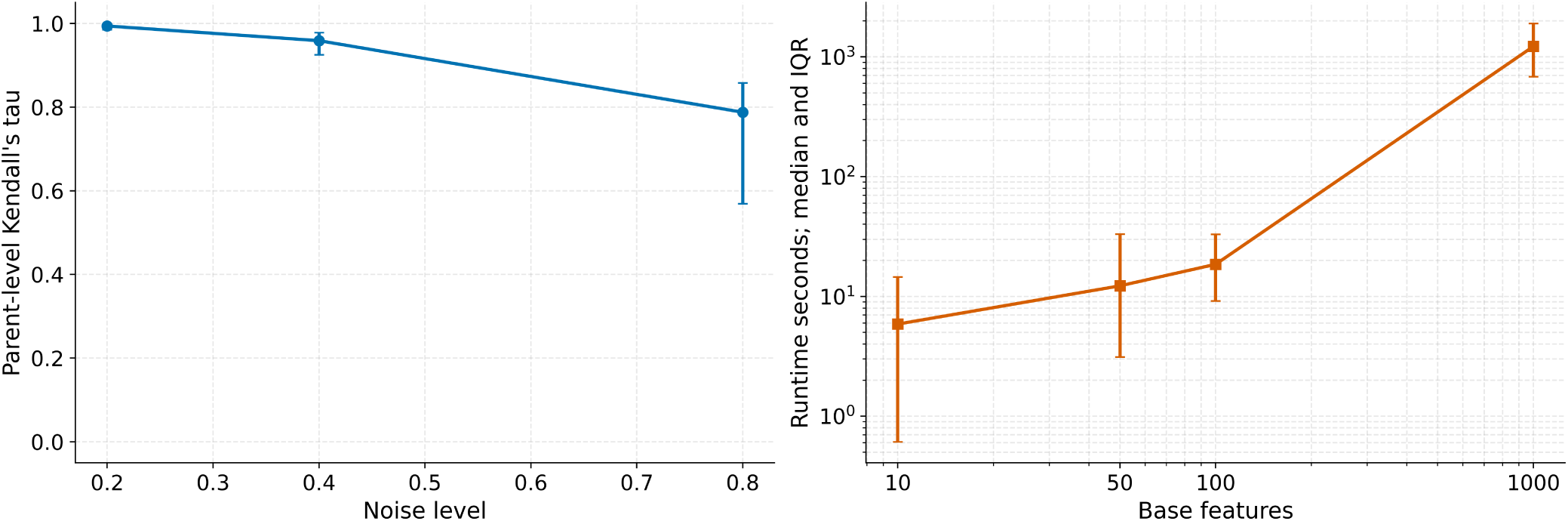
FastEBM performance across the completed correlated-feature synthetic benchmark. As expected, runtime increased with the number of measured features, reflecting the added computational cost of correlated-feature detection and subject-order estimation as dimensionality increased. Parent-level Kendall’s tau accuracy remained high across feature counts but declined with increasing noise, indicating that measurement noise, rather than correlated feature redundancy itself, was the main driver of reduced event-order recovery. Points indicate medians across completed configurations; error bars indicate interquartile ranges.

Accuracy was stable across the larger feature settings. Median Kendall’s tau was 0.933 for 10 base features, 0.950 for 50 base features, 0.964 for 100 base features, and 0.967 for 1,000 base features. Increasing sample size also improved robustness: median tau increased from 0.907 at 250 subjects per group to 0.985 at 2,500 subjects per group, and the high-noise median increased from 0.693 to 0.878 across the same sample-size range (**Table** 5).

**Table 5:** Accuracy and Runtime Comparison Across Different Number of Subjects.

| Number of Subjects | Parent Kendall's tau Median | Runtime Median (seconds) |
| --- | --- | --- |
| 500 | 0.9068 | 4.5061 |
| 1000 | 0.9570 | 10.9142 |
| 2000 | 0.9590 | 16.6586 |
| 5000 | 0.9852 | 65.5463 |

The correlated-feature workflow remained computationally tractable across the completed grid, but runtime increased sharply with feature dimensionality. The median total runtime was 20.45 s (IQR 8.22- 141.73 s) across all configurations. Median runtime increased from 5.88 s for 20 total measured features to 12.26 s for 100 features, 18.53 s for 200 features, and 1,221.91 s for 2,000 features. Runtime also increased with sample size, from a median of 4.51 s at 250 subjects per group to 65.55 s at 2,500 subjects per group. The longest run occurred for 1,000 base features, 2,500 subjects per group, and noise level 0.4, requiring 3,716.45 s. Runtime was driven primarily by correlated-feature detection and first-passage uncertainty/subject-order estimation (**Figure** 6).

### 3.3 TADPOLE

Event sequences generated for the TADPOLE dataset using each EBM method are shown in **Figure** 7. FastEBM’s inferred ordering started with ABETA42 followed by TAU and PTAU, then structural MRI markers (hippocampus and whole brain volume), and finally cognitive scores (ADAS13, MMSE) (**Figure** 8). This ordering is consistent with the hypothetical model of Alzheimer’s disease progression proposed by Jack et al., 2013 and the empirical findings from Bateman et al., 2012 where amyloid-*β* accumulation is the earliest detectable event, followed by tau pathology, brain atrophy, and cognitive decline. In contrast, GMM-EBM and KDE-EBM rank cognitive scores early in the disease progression timeline followed by varying combinations of tau marker abnormality and brain volume loss. Specifically, KDE-EBM ranks tau pathology followed by structural brain atrophy and amyloid-*β* accumulation. For EBM amyloid-*β* accumulation and tau abnormality occur after hippocampus volume decline. DEBM has a similar ranking to GMM-EBM, but it ranks amyloid-*β* abnormality first and total tau last.

**Figure 7:**
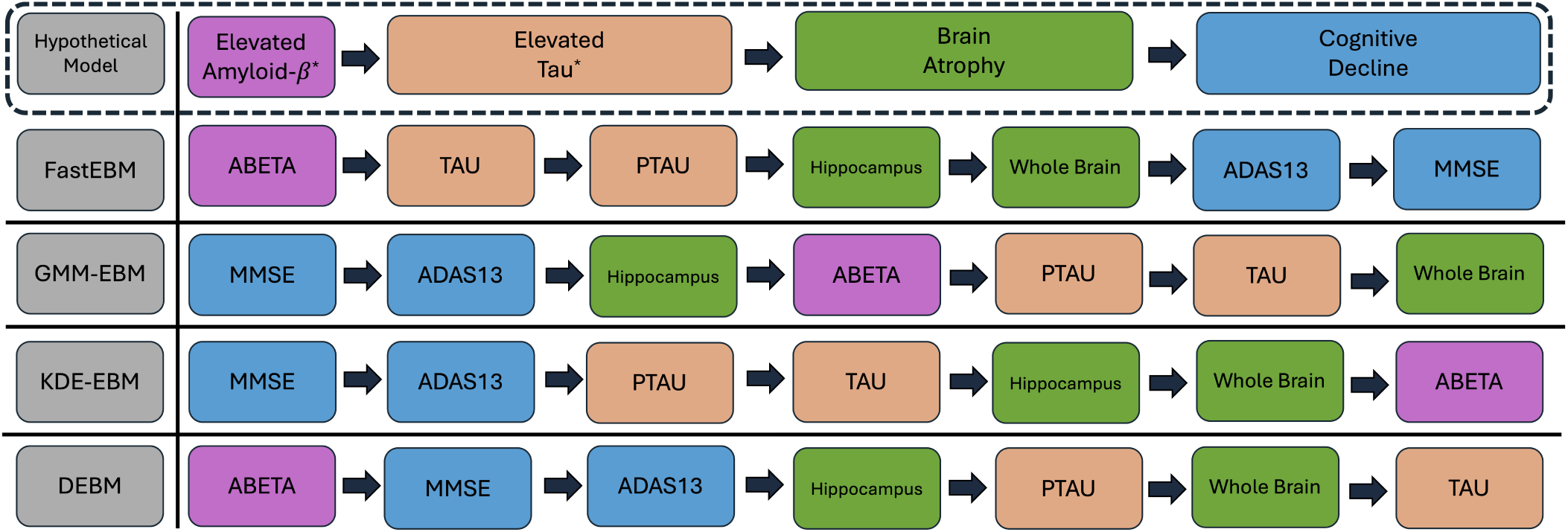
Event ordering estimated from the TADPOLE dataset using FastEBM, GMM-EBM, KDE- EBM, and DEBM, shown relative to the canonical hypothetical model of Alzheimer’s disease progression model (top). *^∗^*For CSF biomarkers, amyloid-*β* levels decrease, whereas tau levels increase.

**Figure 8:**
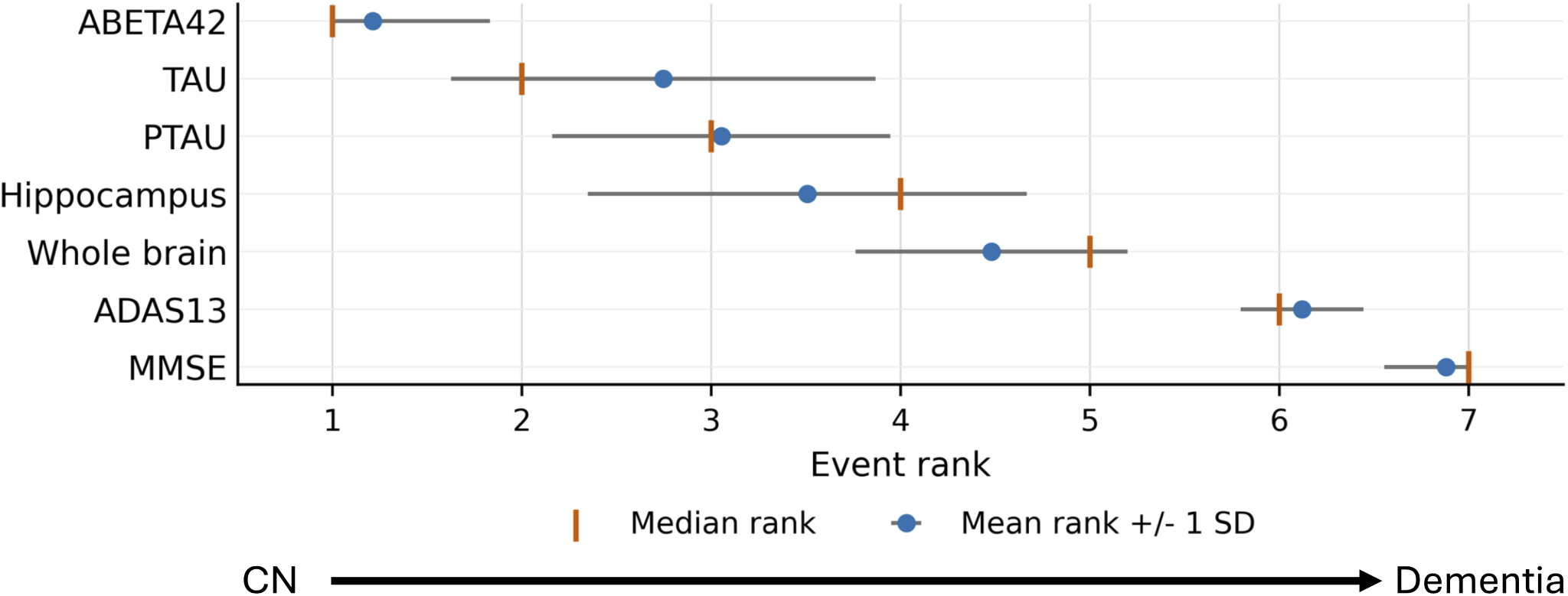
Mean event rank and rank uncertainty for the seven TADPOLE biomarkers. Points show mean rank, horizontal bars show ±1 SD across first-passage uncertainty samples, and orange ticks show median rank.

### 3.4 Regional Alzheimer’s Disease Cortical Tau Progression

FastEBM-derived cortical tau-PET ordering was broadly consistent with Braak-like (Braak & Braak, 1991) tau progression (**Figure** 9). The earliest median event ranks were localized to the entorhinal cortex, followed by a broader temporal-limbic cluster that included inferior/middle temporal, parahippocampal, fusiform, insular, and superior temporal cortices. Subsequent events extended across widespread association cortex, including lateral temporal, frontal, cingulate, parietal, and occipital association regions. The latest events involved primary sensory-motor and visual cortices, including precentral, postcentral, paracentral, cuneus, pericalcarine, and transverse temporal regions.

**Figure 9:**
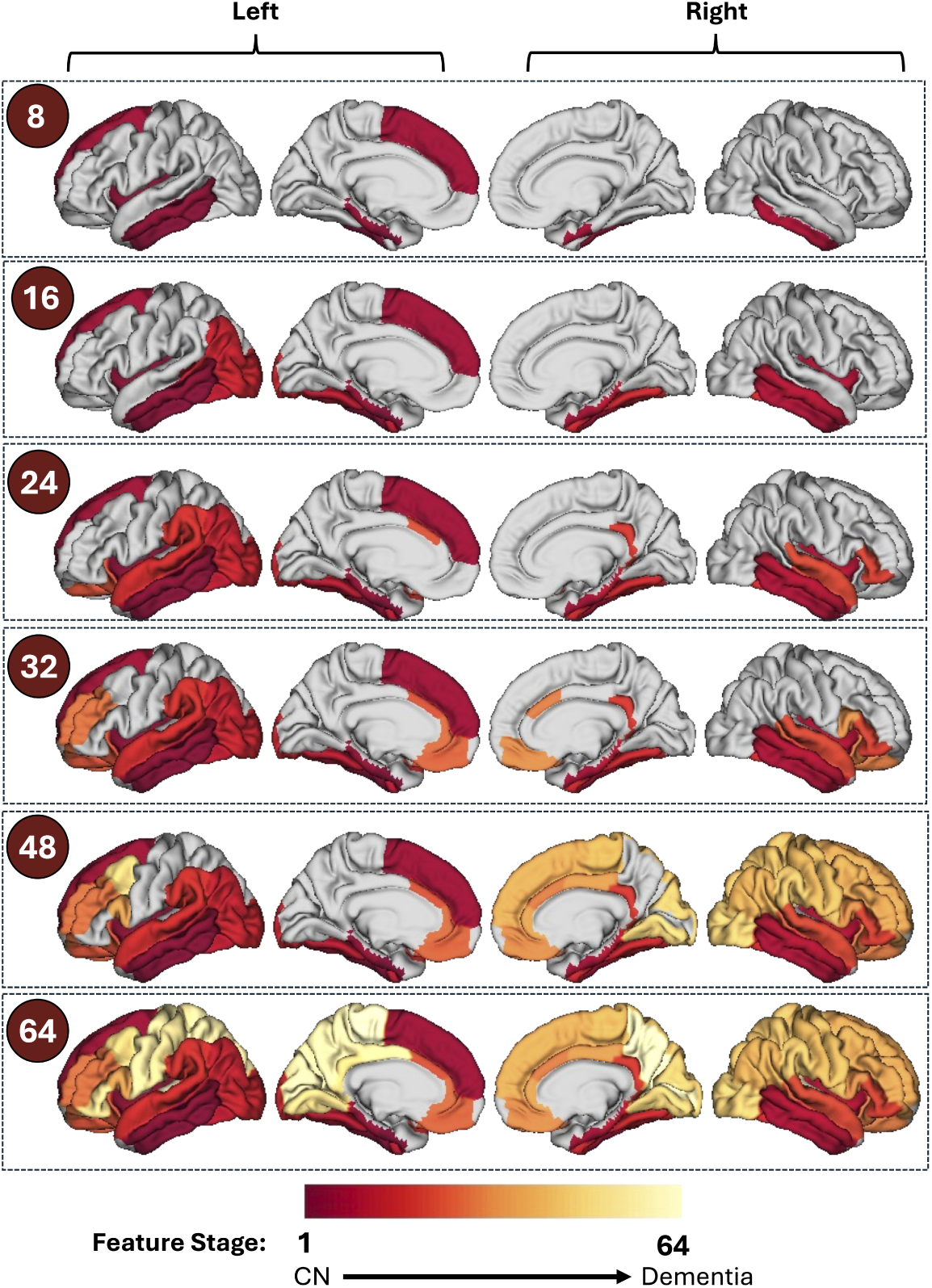
Cortical maps show the cumulative spatial distribution of tau-PET ROIs included by increasing FastEBM event-rank (feature stage) cutoffs. Here, feature stage in the legend is equivalent to the median event rank. At each cutoff, regions with median event rank less than or equal to the displayed threshold are colored, while later-ranked regions remain gray.

The cumulative event-rank cutoff maps suggested a lateralized component to the inferred group-level ordering. Early cutoffs were dominated by left-hemisphere temporal and temporolimbic regions, whereas later cutoffs included a relatively greater proportion of right-hemisphere association regions, followed by bilateral sensorimotor and visual cortices. Event-rank uncertainty is illustrated in **Figure** 10.

**Figure 10:**
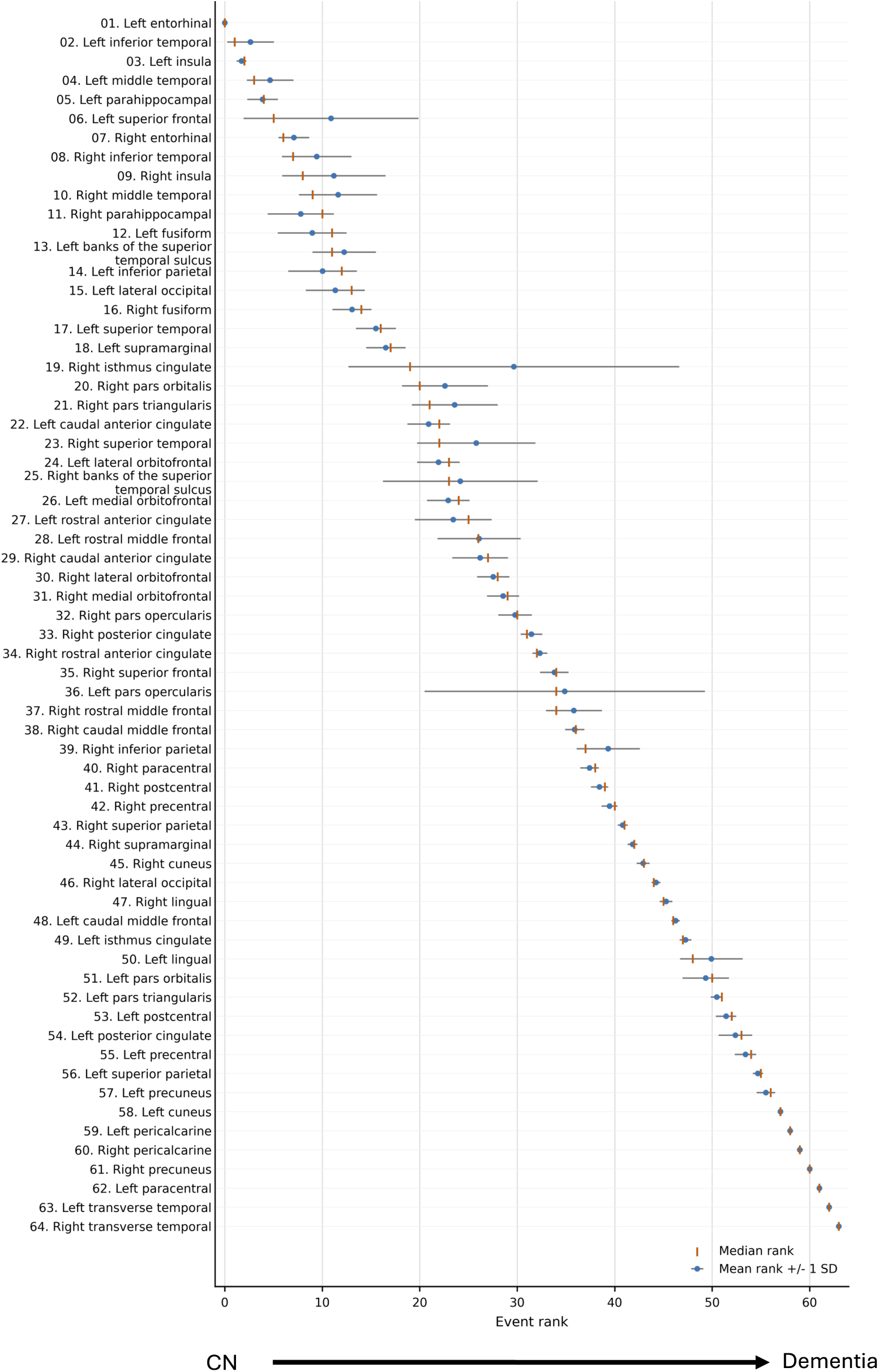
Mean event rank and rank uncertainty for the 64 cortical FreeSurfer tau-PET ROIs. Points show mean rank, horizontal bars show ±1 SD across first-passage uncertainty samples, and orange ticks show median rank.

## 4 Discussion

In this work, we introduced FastEBM, a scalable Markov-chain-based event-based modeling framework that reformulates disease progression inference as a subject-ordering problem on a data-driven manifold. Rather than estimating biomarker event permutations directly through likelihood optimization or sampling, FastEBM models disease progression as a stochastic diffusion process over a subject similarity graph and derives progression structure from the corresponding fundamental matrix. We also proposed an uncertainty-aware extension that uses intrinsic first-passage-time variability to quantify uncertainty in inferred progression ordering. Across synthetic benchmarking experiments and Alzheimer’s disease tau-PET data, FastEBM consistently demonstrated strong event-order recovery accuracy, substantial computational advantages over conventional EBM approaches, and robustness in high-dimensional and correlated-feature settings.

In the performance comparisons with other EBM methods, Deterministic FastEBM and FastEBM consistently produced the highest Kendall’s tau values while remaining faster than the baseline methods. In the main-benchmark configurations, Deterministic FastEBM achieved a median Kendall’s tau of 0.992 with a median runtime of 1.28 s, whereas FastEBM achieved a median tau of 0.997 with a median runtime of 2.23 s. In contrast, DEBM and GMM-EBM required median runtimes on the order of tens to hundreds of seconds while producing substantially lower ordering accuracy. KDE-EBM was both inaccurate and numerically unstable, failing in multiple configurations because covariance matrices were not positive definite. The uncertainty-aware FastEBM extension introduced only a moderate computational overhead relative to the deterministic model. The present results suggest that intrinsic uncertainty estimation based on first-passage-time variability can be integrated into large-scale progression modeling without rendering the method impractical for high-dimensional datasets. At a fixed cohort size, FastEBM’s runtime increased smoothly from fractions of a second to only a few seconds as the number of features increased from 10 to 1,000, while preserving near-perfect event-order recovery. In contrast, DEBM and GMM-EBM exhibited runtime increases spanning several orders of magnitude. The difference was especially noticeable for DEBM, whose runtime reached approximately 2.36 × 10^5^ s in the largest feature configuration examined. Current event-based models scale poorly because they rely on discrete combinatorial optimization. When framed as decision problems, these tasks are NP-hard, meaning no exact, polynomial-time algorithms exist to solve them (Schrijver, 2003). Consequently, as the number of features increases, the number of possible permutations grows factorially (*p*!). This massive search space makes random search algorithms, such as Markov Chain Monte Carlo (MCMC), highly prone to long run- times and suboptimal results. Similar behavior was observed during subject-scaling experiments, where FastEBM maintained stable accuracy while runtime growth remained comparatively modest. These findings suggest that replacing combinatorial event-permutation inferences with Markov diffusion geometry substantially alters the computational scaling properties of event-based modeling.

The high-dimensional low-sample size stress tests are particularly relevant because many neuroimaging and biomedical datasets naturally operate in this regime (Mwangi et al., 2014). Conventional EBM approaches degraded substantially under these conditions, with large reductions in ordering accuracy and frequent numerical instability. In contrast, FastEBM retained strong event-order recovery even under severe high-dimensional low-sample size conditions. Deterministic FastEBM completed all stress-test configurations with a median Kendall’s tau of 0.965 and a median runtime of only 0.034 s, while FastEBM achieved comparable accuracy with runtime under 5 s. These results indicate that the proposed Markov-chain representation is particularly well suited to the geometric structure of high-dimensional low-sample size progression problems. Noise level was the dominant determinant of performance across all experiments. At low and moderate noise levels, both FastEBM variants recovered event orderings with near-perfect accuracy across most synthetic configurations. At the highest noise level, all methods deteriorated, but the FastEBM approaches retained substantially more event-order information than the alternatives. Even at noise level 0.8, Deterministic FastEBM maintained median Kendall’s tau values above 0.9 in the main benchmark and approximately 0.75 in the high-dimensional low-sample size stress tests. This suggests that the diffusion-based subject representation captures global progression geometry in a manner that is comparatively robust to local measurement noise.

To assess FastEBM’s sensitivity to correlated and redundant biomarkers, we performed synthetic dataset experiments with correlated features generated from parent features using FastEBM with the uncertainty workflow. These experiments demonstrated that FastEBM remained robust despite substantial redundancy. Parent-level event-order recovery remained high across nearly all completed configurations, with overall median Kendall’s tau 0.957. Accuracy remained stable across larger feature settings, indicating that the framework successfully mitigated much of the instability introduced by correlated duplicate biomarkers. As in Experiment 1, noise level remained the primary determinant of performance. The smallest feature setting showed the greatest sensitivity to severe noise, whereas the larger feature settings retained strong ordering accuracy even in challenging conditions. This behavior suggests that the progression geometry inferred by FastEBM becomes increasingly stable as more informative biomarkers are incorporated, provided redundancy is appropriately controlled. The runtime behavior of the correlated-feature workflow is also informative. Runtime increased substantially at the largest dimensionalities, particularly for the 2,000-feature configurations, indicating that correlated-feature detection and uncertainty estimation remain important computational bottlenecks. Nevertheless, the framework remained tractable across the completed experimental grid.

The uncertainty-aware extension represents a conceptual departure from conventional EBM uncertainty estimation strategies. Classical approaches typically estimate uncertainty using bootstrap resampling, Monte Carlo permutation sampling, or posterior sampling over event sequences (Firth et al., 2020; Fonteijn et al., 2012; Young et al., 2014). In contrast, FastEBM derives uncertainty directly from intrinsic stochastic variability of the inferred Markov process. First-passage-time variance quantifies fluctuations in the same diffusion geometry used to estimate subject ordering and disease progression scores. Consequently, uncertainty is represented as variability within the inferred progression manifold itself rather than solely as instability under repeated cohort resampling. This distinction is important because it links uncertainty directly to progression geometry and transition structure. The Markovian uncertainty is conditional on the observed cohort and the fitted transition graph. Thus, the resulting intervals represent a model-internal propagation sensitivity rather than a bootstrap- or posterior-style uncertainty estimate over all sources of sampling, measurement, and model uncertainty. The resulting probabilistic ordering framework allows identification of stable versus ambiguous regions of progression while remaining computationally efficient relative to large-scale sampling procedures.

Although the absolute ground truth ordering for Alzheimer’s disease progression in a specific set of participants may be unknown, using the multimodal Alzheimer’s disease TADPOLE biomarker dataset, FastEBM identified a progression sequence consistent with canonical Alzheimer’s disease models, with amyloid-*β* abnormalities preceding tau pathology, followed by structural brain atrophy and subsequent cognitive decline (Bateman et al., 2012; Jack et al., 2013). Previous EBM studies using TADPOLE have reported alternative biomarker orderings (Du et al., 2023; Venkatraghavan et al., 2019).

FastEBM’s application to estimate the regional spread of cortical tau-PET across Alzheimer’s disease progression showed that FastEBM recovered biologically plausible progression trajectories consistent with established models of Alzheimer’s disease tau spread. The pattern indicated tau spreading from medial temporal and temporal-limbic regions toward the broader association cortex and, finally, primary sensory-motor and visual cortices (Mormino & Papp, 2018). However, the regional ordering also showed deviations from this coarse staging pattern. These deviations may reflect known heterogeneity in AD tau topography, as growing evidence suggests that AD tau spread is not restricted to a single Braak-like trajectory. Neuropathological studies have identified distinct AD subtypes, including typical, limbic-predominant, and hippocampal-sparing patterns (Murray et al., 2011; Whitwell et al., 2012). More recently, tau-PET studies have shown that tau deposition can follow different spatiotemporal trajectories, including medial-temporal-sparing, posterior, and lateral temporal patterns, as well as asymmetric left- or right-hemisphere involvement (Ossenkoppele et al., 2016; Vogel et al., 2021). In this context, the lateralized pattern observed here may reflect cohort heterogeneity, where asymmetric tau-spread patterns across individuals contribute to a group-level ordering with earlier left temporal involvement and a relative shift toward right-hemisphere association regions at later event ranks. The high regional resolution of FastEBM’s ordering may therefore expose lateralized or regionally specific deviations from coarse Braak composite staging that would otherwise be averaged out.

Unlike previous EBM studies that have relied on a relatively small number of features (Fonteijn et al., 2012; Young et al., 2014), FastEBM is designed to significantly increase spatial resolution. Although the present study considered 64 cortical regions of interest, the proposed framework is not inherently limited to this level of feature size as demonstrated using simulated data. The computational efficiency of FastEBM allows for extensions to increasingly finer spatial representations, including vertex-wise cortical measurements, to enable more detailed characterization of regional disease progression.

Finally, we note two main limitations of our framework. First, although the synthetic experiments covered a broad range of dimensionalities, cohort sizes, and noise conditions, the simulations do not fully capture the complexity of real disease heterogeneity, subtype structure, or nonlinear biomarker trajectories. FastEBM estimates a single dominant event ordering across all subjects assuming a single subtype. Therefore, the inferred ordering may reflect an average or mixture of subtype-specific trajectories. Further, each feature is assigned a single dominant change point along the inferred disease-progression axis, which may be less robust for features with different trajectories than sigmoidal or monotonic. Future work will evaluate the sensitivity of the change point procedure across a broader family of trajectory shapes and apply FastEBM within empirically defined or data-driven subtypes. Second, the correlated-feature workflow currently relies on heuristic grouping and dimensionality-reduction procedures to identify and merge redundant features before model fitting. Although this approach reduced redundancy in the present experiments, the grouping step introduces an additional modeling decision that can influence the inferred subject ordering and downstream event ranks. Future work will evaluate more model-based approaches for handling correlated features, such as sparse representation learning or latent-variable models.

In summary, FastEBM provides a scalable and robust alternative to conventional event-based modeling approaches. Across synthetic and real-data experiments, the framework consistently preserved strong event-order recovery accuracy while dramatically reducing computational cost relative to existing EBM implementations. The uncertainty-aware extension further enables probabilistic characterization of disease progression directly from intrinsic Markov-chain variability while remaining computationally practical. Together, these results suggest that diffusion-based Markov geometry provides a promising foundation for scalable, uncertainty-aware disease progression modeling in high-dimensional biomedical datasets. An open-source implementation of FastEBM is available at https://github.com/sjusc07/FastEBM.

## Data and Code Availability

FastEBM is an open-source package. The code may be read in full at: https://github.com/sjusc07/FastEBM.

## Acknowledgements

This work was supported by grant R01AG087513 from the National Institute on Aging at the National Institutes of Health, by grant S10OD032285 from the National Institutes of Health, by grant AARG-23- 1149996 from US Alzheimer’s Association.

We thank Dr. Esther E. Bron and Laura Monteiro Rente Dias (Erasmus MC, Rotterdam, The Netherlands) for valuable discussions and for inspiring the high-dimensional low-sample size stress tests, which provided important insights into the behavior of the model.

Data used in the preparation of this article were obtained from the Alzheimer’s Disease Neuroimaging Initiative (ADNI) database (adni.loni.usc.edu). The ADNI was launched in 2003 as a public-private partnership, led by Principal Investigator Michael W. Weiner, MD. The primary goal of ADNI has been to test whether serial magnetic resonance imaging (MRI), positron emission tomography (PET), other biological markers, and clinical and neuropsychological assessment can be combined to measure the progression of mild cognitive impairment (MCI) and early Alzheimer’s disease (AD). For up-to-date information, see www.adni-info.org. Data collection and sharing for the Alzheimer’s Disease Neuroimaging Initiative (ADNI) is funded by the National Institute on Aging (National Institutes of Health Grant U19 AG024904). The grantee organization is the Northern California Institute for Research and Education.

In the past, ADNI has also received funding from the National Institute of Biomedical Imaging and Bioengineering, the Canadian Institutes of Health Research, and private sector contributions through the Foundation for the National Institutes of Health (FNIH) including generous contributions from the following: AbbVie, Alzheimer’s Association; Alzheimer’s Drug Discovery Foundation; Araclon Biotech; BioClinica, Inc.; Biogen; Bristol-Myers Squibb Company; CereSpir, Inc.; Cogstate; Eisai Inc.; Elan Pharmaceuticals, Inc.; Eli Lilly and Company; EuroImmun; F. Hoffmann-La Roche Ltd and its affiliated company Genentech, Inc.; Fujirebio; GE Healthcare; IXICO Ltd.; Janssen Alzheimer Immunotherapy Research & Development, LLC.; Johnson & Johnson Pharmaceutical Research &Development LLC.; Lumosity; Lundbeck; Merck & Co., Inc.; Meso Scale Diagnostics, LLC.; NeuroRx Research; Neurotrack Technologies; Novartis Pharmaceuticals Corporation; Pfizer Inc.; Piramal Imaging; Servier; Takeda Pharmaceutical Company; and Transition Therapeutics.

